# Purification of functional *Plasmodium falciparum* tubulin allows for the identification of parasite-specific microtubule inhibitors

**DOI:** 10.1101/2021.05.25.445550

**Authors:** William Graham Hirst, Dominik Fachet, Benno Kuropka, Christoph Weise, Kevin Saliba, Simone Reber

## Abstract

Cytoskeletal proteins are essential for parasite proliferation, growth, and transmission, and therefore represent promising drug targets. While αβ-tubulin, the molecular building block of microtubules, is an established drug target in a variety of cancers, we still lack substantial knowledge of the biochemistry of parasite tubulins, which would allow us to exploit the structural divergence between parasite and human tubulins. Indeed, mechanistic insights have been limited by the lack of purified, functional parasite tubulin. In this study, we isolated *Plasmodium falciparum* tubulin that is assembly-competent and shows specific microtubule dynamics *in vitro*. We further present mechanistic evidence that two compounds selectively interact with parasite over host microtubules and inhibit *Plasmodium* microtubule polymerization at substoichiometric compound concentrations. The ability of compounds to selectively disrupt protozoan microtubule growth without affecting human microtubules provides the exciting possibility for the targeted development of novel antimalarials.

## INTRODUCTION

Microtubules and their molecular building block αβ-tubulin have gained outstanding importance as targets for drug development. Disrupting microtubule dynamics affects cytoskeletal function, interphase cell signaling, and cell division. Colchicine, paclitaxel, and vinca alkaloids are the earliest plant-derived microtubule-targeting agents and are broadly classified into two categories: microtubule stabilizing agents - they promote tubulin polymerization and stabilize microtubules against depolymerization - and microtubule destabilizing agents, which depolymerize microtubules and/or prevent *de novo* microtubule formation (reviewed by Florian and Mitchison, 2016, Steinmetz and Prota 2018). Many tubulin-binding drugs are already used as therapeutics or are in clinical trials. Predominant among these small molecules are paclitaxel (Taxol®) and docetaxel (Taxotere®), which have demonstrated their activity against a variety of malignancies (Florian and Mitchison, 2016). The effectiveness of microtubule-targeting drugs, however, is limited by their toxicity. Above all, they cause neutropenia and neurotoxicity, the latter likely by disruption of microtubule-mediated axonal transport in neurons (reviewed by Serpico et al. 2020).

As other eukaryotic cells, *Plasmodium* species rely on microtubules to build their cytoskeleton, flagella, and the mitotic spindle, making them a prime drug target. Although there is a high degree of amino-acid conservation between human and *P. falciparum* α- and β-tubulin, sharing approximately 83.7 and 88.5 % identity respectively, *Plasmodium* tubulin is more similar to plant tubulin than to mammalian tubulin. This opens up the possibility of identifying compounds that selectively disrupt parasite microtubules but do not affect the host cell cytoskeleton. In addition, taxanes have been shown to be efficient against single and multidrug resistant parasites, implying that tubulin-binding drugs may be used to overcome the problem of multidrug resistance (Schrevel et al., 1994, Dieckmann-Schuppert and Franklin, 1989). Although a number of microtubule inhibitors have been explored in *Plasmodium* (reviewed by Kappes and Rohrbach 2007), our molecular and biochemical knowledge of *P. falciparum* tubulin and microtubules is limited. For example, it remains to be shown whether compounds of interest specifically target parasite microtubules without affecting the host cell cytoskeleton. Such mechanistic insights have so far been limited by the lack of functional parasite tubulin.

The interest in the identification of parasite-specific microtubule-binding drugs is demonstrated in a variety of studies, including those that use homology modeling and molecular docking (Lyons-Abbott et al. 2010, Chakrabarti et al. 2013, Morrissette 2015, Soleilhac et al. 2018). The limitation of most of these studies, however, is the use of bovine or porcine tubulin structures as templates, as no parasite tubulin structure is available. Indeed, during the last 50 years, most biochemical studies investigating microtubule dynamics and microtubule-targeting agents have used tubulin purified from mammalian brain tissue, typically bovine or porcine brain (Gell et al., 2011), with the consequence of studying heterologous systems. In this study, we report the purification and characterization of *P. falciparum* tubulin from infected red blood cells. Importantly, the highly purified tubulin is fully functional as it efficiently assembles into microtubules that show specific parameters of dynamic instability. We further show for two compounds selective toxicity for parasite, but not for human, tubulin. Taken together, these microtubule-binding drugs might therefore represent scaffolds that act as leads for the design of new-generation antimalarials targeting microtubules.

## RESULTS

### Purification of assembly-competent tubulin from *P. falciparum*-infected erythrocyte cultures

As the asexual blood stages of *P. falciparum* are responsible for the clinical symptoms of malaria and can be readily cultured in the lab (Trager and Jensen 1976, Maier et al. 2018), we decided to purify tubulin from infected red blood cells (Fig. 1a). By sequentially lysing first the red blood cells and, in a second step, the parasites, we were able to separate the cytoplasmic content of the host cell from that of the parasites (Supplementary Fig. 1a). In the following step, αβ-tubulin was purified by affinity chromatography (Widlund et al. 2011, Reusch et al. 2020) from the parasite lysate (Fig. 1b). As some previous reports suggested that human erythrocytes contained tubulin (Bryk and Wiśniewski 2017, Nigra et al. 2020), we monitored the presence of tubulin throughout the sequential host and parasite cell lysis (Supplementary Fig. 1b and c) and concluded that host tubulin is absent from isolated parasite lysates (see also Fig. 2a) and purified tubulin. The protocol we applied captured the full complement of *P. falciparum* tubulin, indicated by a complete depletion of tubulin from the lysate (Fig. 1c, F= flowthrough), confirming that we did not enrich for a tubulin subpopulation (e.g. a specific isoform or post-translational modification). This means that the purified tubulin represents the physiological tubulin proteome. Importantly, and in contrast to previous attempts (Dempsey et al. 2013, Fennell et al. 2006), our purified *P. falciparum* tubulin was assembly-competent and showed characteristic microtubule dynamic instability as visualized by total internal reflection fluorescence (TIRF) microscopy (Fig. 1d and e, Supplementary movie S1). To our knowledge, this is the first time that active, assembly-competent tubulin has been purified from *P. falciparum*.

**Figure 1.**
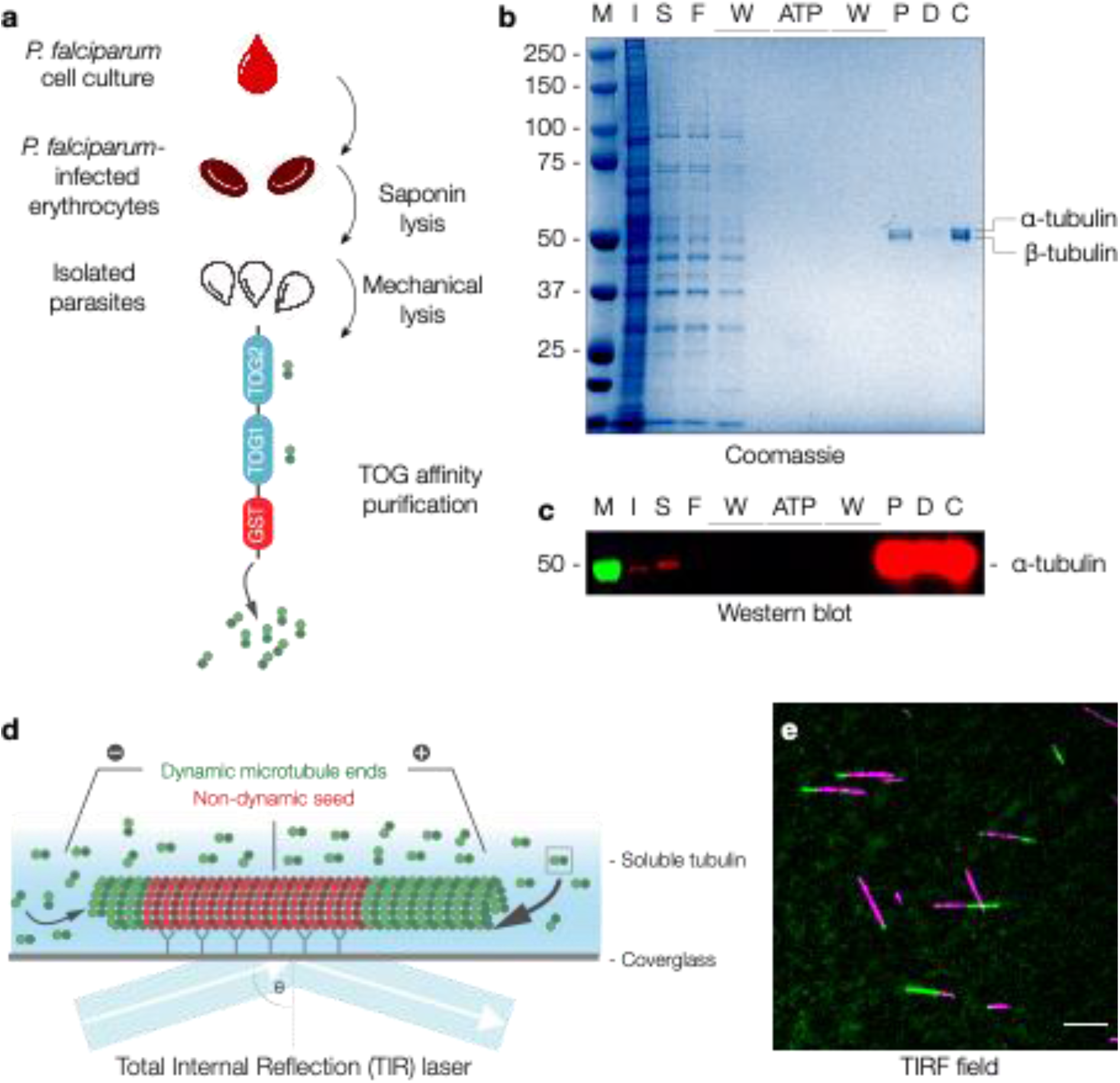
Affinity purification of blood-stage *P. falciparum* tubulin. (**a**) Strategy detailing the purification of *P. falciparum* tubulin. (**b**) Coomassie-stained SDS-PAGE of the individual purification steps from *P. falciparum* lysate. M: Marker, I: Input, S: Supernatant, F: Flowthrough, W: Wash, ATP: ATP wash, P: Peak fraction, D: Desalt, C: Concentrated. (**c**) Western blot analysis with anti-α-tubulin antibody of samples shown in (**b**). (**d**) Schematic of a microtubule assembly assay visualized by total internal reflection fluorescence (TIRF) microscopy using GMPCPP-stabilized microtubule seeds (red) as nucleation templates for dynamic microtubules (green). (**e**) Representative TIRF microscopy image of dynamic *P. falciparum* microtubules (green) grown from stabilized seeds (magenta) at 37°C at 6 μM tubulin. Scale bar: 5 μm. See also Supplementary movie S1.

**Figure 2.**
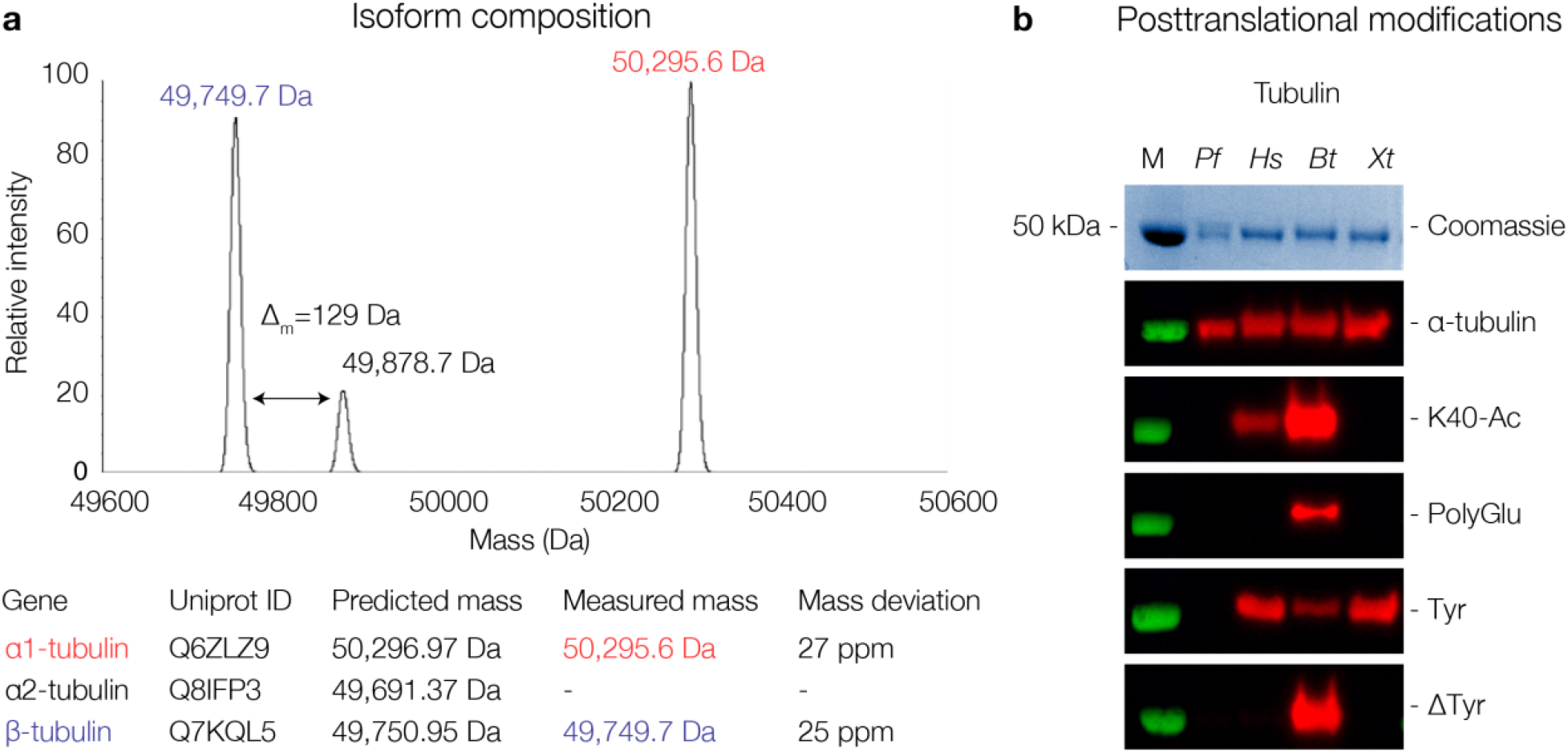
Isoform composition and posttranslational modifications of purified blood-stage *P. falciparum* tubulin. (**a**) Deconvoluted mass spectrum of purified *P. falciparum* tubulin. Individual tubulin isoforms are labeled in the spectrum and measured masses are compared with predicted masses in the table below. The raw mass spectrum used for deconvolution is shown in Supplementary Fig. 2C (**b**) Western blots probing posttranslational modifications found in purified *P. falciparum* (*Pf*), HEK cells (*Hs*), bovine brain (*Bt*), and *Xenopus laevis* (*Xl*) tubulin. α-tubulin is a loading control, K40-Ac recognizes acetylated lysine at position 40 of α-tubulin, Poly-Glu recognizes epitopes containing acidic residues modified with a chain of at least 2 glutamyl residues, Tyr recognizes the C-terminal EEY epitope of tyrosinated tubulin, ΔTyr recognizes the detyrosinated C-terminus of α-tubulin.

### Blood-stage *P. falciparum* tubulin is composed of the α1- and β-isoform

Purified tubulin populations are usually a complex mixture of isoforms with various posttranslational modifications specific for a given cell line or species (Chaaban et al. 2018, Hirst et al. 2020a). The *P. falciparum* genome, however, only encodes two α-tubulin and one β-tubulin isoforms (Supplementary Fig. 2a), and antibody-based experiments suggest that α1-tubulin is the predominant isoform at the protein level during blood stages (Delves et al. 1989, Holloway 1989, Holloway 1990, Rawlings 1992). Consistently, expression data show higher expression of α1-tubulin versus α2-tubulin during the blood stages (Toenhake et al. 2018, Josling et al. 2015). To characterize the isoform composition present in our purified *P. falciparum* tubulin, we digested the tubulin and analyzed it by matrix-assisted laser desorption/ionization mass spectrometry (MALDI-MS). The corresponding peptide mass fingerprints revealed the presence of *P. falciparum* α1- and β-tubulin (Supplementary Fig. 2b) but not the α2-isoform. In addition, no other significant peptides were identified. This further confirmed that purified *P. falciparum* tubulin was free of human host tubulin and free of potentially contaminating microtubule-associated proteins (MAPs) and motors. As a complementary approach, intact proteins were analyzed by liquid chromatography electrospray ionization - mass spectrometry (LC-ESI-MS) (Fig. 2a and Supplementary Fig. 2c), which showed a main peak at 50,296 Da that matches the mass predicted for α1-tubulin (Uniprot ID Q6ZLZ9, 50,297 Da), and a second major peak at 49,750 Da that corresponds to β-tubulin (Uniprot ID Q7KQL5, 49,751 Da). In addition, a smaller peak was detected at 49,879 Da, which is likely explained by the addition of a single glutamyl modification (Δ_m_ = +129 Da) on β-tubulin. Notably, a peak that would correspond to α2-tubulin at 49,691 Da (Uniprot ID Q8IFP3) is absent from the spectrum. In addition to genetic diversity, tubulins are functionalized with diverse posttranslational modifications (PTMs), such as acetylation, polyglutamylation, phosphorylation, and (de-)tyrosination (as reviewed in Gadadhar et al. 2017, Roll-Mecak 2019). Although it has been shown that *P. falciparum* α1-tubulin can be acetylated *in vitro* (Read et al. 1993), we find no detectable acetylation neither by mass spectrometry nor western blot analysis (Fig. 2a and b, K-40 Ac). Recent studies using expansion microscopy on *P. falciparum* schizonts have shown that hemispindles are not polyglutamylated, whereas the subpellicular microtubules were polyglutamylated (Bertiaux et al. 2020). Our whole-protein mass spectrometry data found the purified tubulin to be monoglutamylated (Figure 2a), giving a potential explanation for why the tubulin is not recognized by an antibody against polyglutamylation (Figure 2b, Poly-Glu, see discussion). Together, these results from the two mass spectrometry approaches and western blot analysis suggest that α1-tubulin is the predominant α-tubulin isoform of blood-stage *P. falciparum* together with β-tubulin, with only minor posttranslational modifications.

### Microtubules assembled from purified *P. falciparum* tubulin have similar dynamic characteristics to mammalian microtubules

In the *P. falciparum* blood stages, microtubules are found as both stable subpellicular microtubules and dynamic spindle microtubules (Bannister et al. 2003, Fennell et al. 2006). However, the intrinsic dynamic properties of *P. falciparum* microtubules have not been characterized to date. Using an *in vitro* reconstitution assay together with TIRF microscopy, we assessed the dynamic parameters of purified *P. falciparum* tubulin in comparison to tubulin purified from bovine brain. Kymographs of dynamic microtubules showed that *P. falciparum* nucleated from GMPCPP-templates at tubulin concentrations as low as 6 μM (Figure 1e) while the nucleation threshold for brain tubulin is at around 9 μM (Wieczorek et al. 2014). Quantifying and comparing parameters of microtubule dynamic instability at 9 μM tubulin and 37°C (Fig. 3a), we found *P. falciparum* microtubules to have a polymerization velocity of 0.438±0.099 μm/min (712±161 tubulin dimers/s), which is comparable to bovine microtubules (0.417±0.108 μm/min, 678±176 tubulin dimers/s, Fig. 3b, n = 106 and 318, respectively, p<0.05, g=0.20). *Plasmodium* microtubules, however, depolymerize significantly faster than bovine microtubules (13.35±6.42 μm/min, 21700±10400 tubulin dimers/s versus 7.78±3.52 μm/min, 12600±5720 dimers/s, Fig. 3c, n = 96 and 174, respectively, p<0.0001, g=1.17). The catastrophe frequency (Fig. 3d), reported as the inverse of microtubule lifetime, was 0.0055±0.0034 s^−1^ versus 0.0076±0.0047 s^−1^ (n = 106 and 318, respectively, p<0.0001, g=0.49) and the rescue frequency 0.0058±0.0218 s^−1^ versus 0.0472±0.0637 s^−1^ (Fig. 3e, n = 96 and 174, respectively, p<0.0001, g=0.78). Taken together, our results demonstrate that *P. falciparum* microtubules display parameters of dynamic instability, with a similar growth velocity but a significantly increased depolymerization velocity and decreased catastrophe and rescue frequencies when compared to the *in vitro* dynamics of bovine brain microtubules.

**Figure 3.**
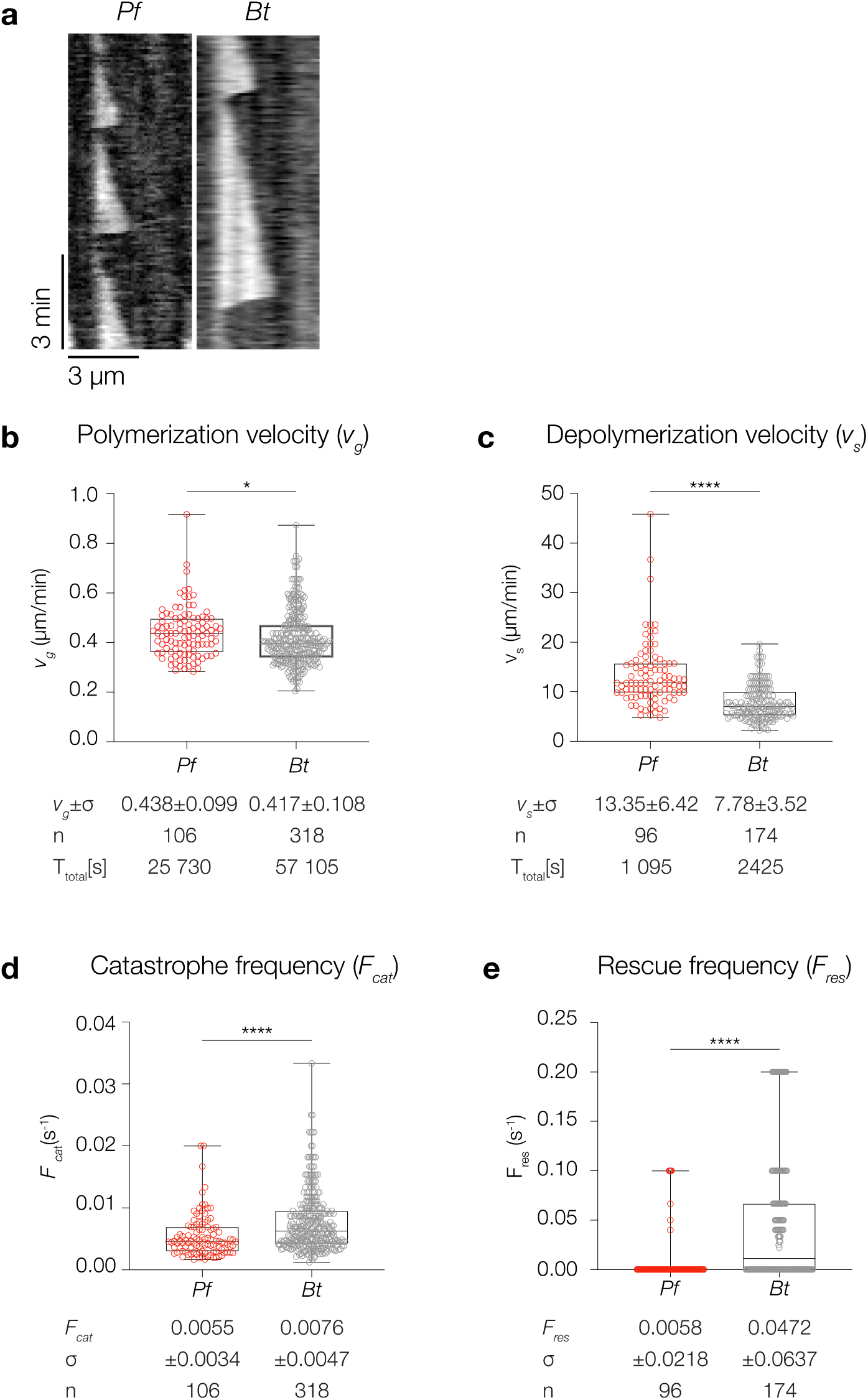
*P. falciparum* microtubules exhibit dynamic properties similar to mammalian microtubules *in vitro*. (**a**) Representative TIRF kymographs of dynamic *P. falciparum* and bovine microtubules at 37°C and 9 μM tubulin. (**b-e**) Parameters of dynamic instability measured at 37°C and 9 μM tubulin. All values were obtained from measurements of microtubules pooled over at least three independent experiments and all p-values are calculated with the Mann-Whitney test. For the modified box-and-whiskers plots the boxes range from 25th to 75th percentile, the whiskers span the range, and the horizontal line marks the median value. (**b**) *Pf* microtubules grow at 0.438±0.099 μm/min (712±161 dimers/min), *Bt* microtubules at 0.417±0.108 μm/min (678±176 dimers/min) with p (*Pf*, *Bt*) <0.05, g=0.20. (**c**) *Pf* microtubules depolymerize at 13.35±6.42 μm/min (21700± 10400 dimers/min), *Bt* microtubules at 7.78±3.52 μm/min (12600±5720 dimers/min) with p (*Pf*, *Bt*) <0.0001, g=1.17. (**d**) Catastrophe frequencies are reported as the inverse of microtubule lifetimes. *Pf* microtubules catastrophe at 0.0055±0.0034 s^−1^, *Bt* microtubules at 0.0076±0.0047 s^−1^ with p (*Pf*, *Bt*) <0.0001, g=0.49. (**e**) Rescue frequencies are reported as the inverse of the duration of each depolymerization event. Events without a rescue are given a value of zero. *Pf* microtubules rescue at 0.0058±0.0218 s^−1^, *Bt* microtubules at 0.0472±0.0637 s^−1^, with p (*Pf*, *Bt*) <0.0001, g=0.78. Total time (T_total_) of observed microtubule growth and shrinkage and number of events (n) indicated.

### Human tubulin purified from HEK293 cells is a permissible comparative tubulin to characterize parasite-specific microtubule inhibitors

As our aim was to characterize parasite-specific microtubule inhibitors that do not affect host microtubules, we first aimed to identify a human tubulin population derived from actively dividing cells with dynamic properties comparable to those of *P. falciparum* microtubules, the rationale being that microtubules with similar dynamics will allow for robust interpretation of inhibition experiments. Indeed, tubulin purified from the human kidney cell line HEK293 (Supplementary Fig. 3a) showed similar microtubule dynamics when compared to *P. falciparum* microtubules (Fig. 4a and b). At 6 μM tubulin, *P. falciparum* microtubules exhibited a slightly higher polymerization velocity (0.385±0.097 μm/min vs. 0.349±0.074 μm/min, p<0.0001, g=0.42, Fig. 4c), a similar depolymerization velocity (6.62±3.36 s^−1^ versus 5.46±2.22 s^−1^, p=0.0002, g=0.40, Fig. 4d), a comparable catastrophe frequency (0.0047±0.0035 s^−1^ vs 0.0058±0.0012 s^−1^, p<0.0001, g=0.42, Fig. 4e) and a lower rescue frequency (0.0120±0.0251 s^−1^ vs 0.0494±0.0284 s^−1^, p<0.0001, g=1.40, Fig. 4f). Therefore, we conclude that human tubulin purified from actively dividing HEK293 cells is a suitable benchmark for the identification and characterization of parasite-specific microtubule inhibitors as both microtubule species display comparable dynamics in *in vitro* reconstitution assays.

**Figure 4.**
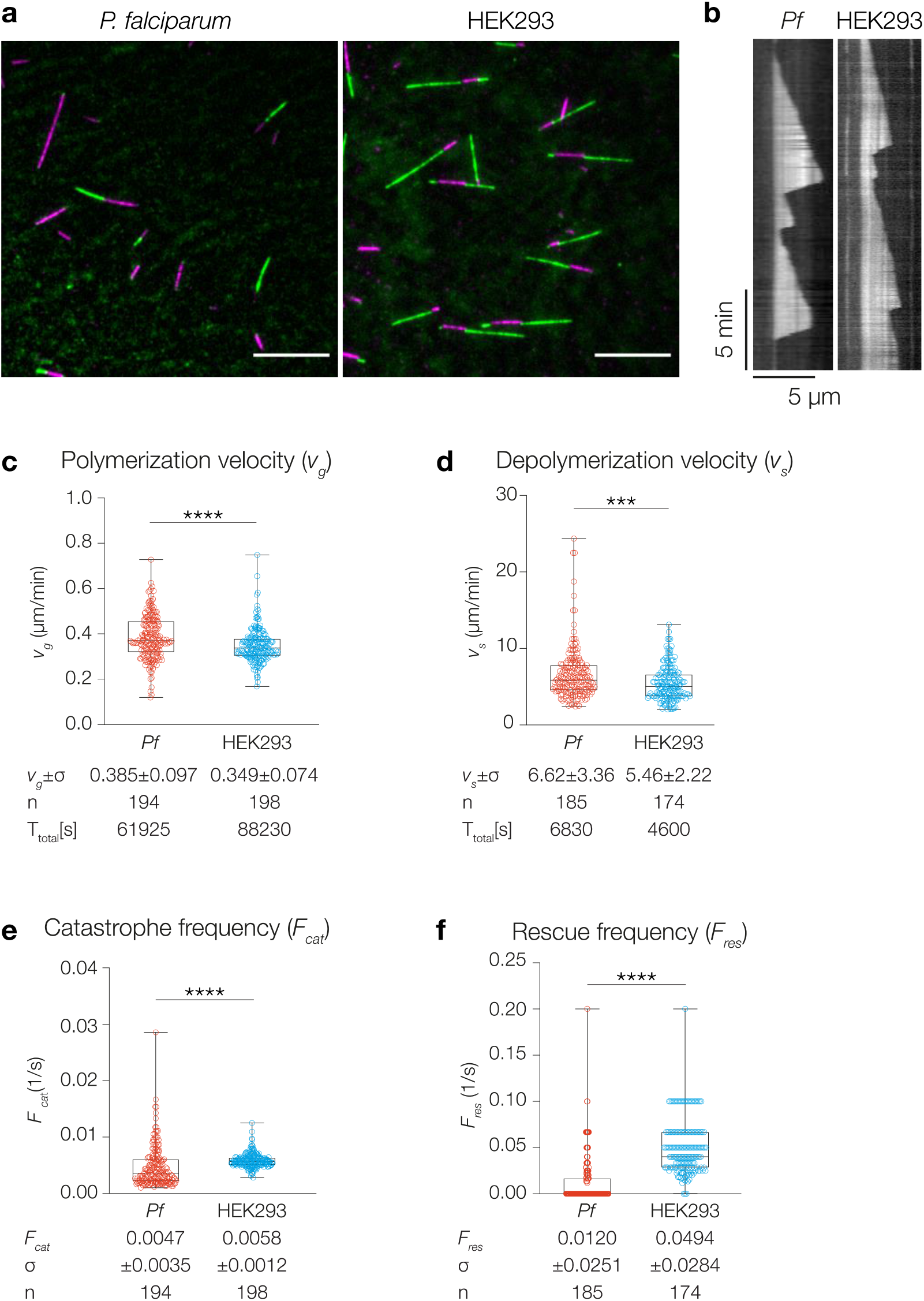
Human tubulin purified from HEK293 cells is a permissible comparative tubulin to characterize parasite-specific microtubule inhibitors. (**a**) Representative TIRF microscopy image of dynamic *P. falciparum* and human microtubules (green) grown from stabilized seeds (magenta) at 37°C and 6 μM tubulin. Scale bar: 5 μm. (**b**) Representative TIRF kymographs of dynamic *P. falciparum* and HEK293 microtubules. (**c-f**) Parameters of dynamic instability measured at 37°C and 6 μM tubulin. All values were obtained from measurements of microtubules pooled over at least three independent experiments and all p-values are calculated with the Mann-Whitney test. For the modified box-and-whiskers plots the boxes range from 25th to 75th percentile, the whiskers span the range, and the horizontal line marks the median value. (**c**) *Pf* microtubules grow at 0.385±0.097 μm/min (626±158 dimers/min), *Hs* microtubules at 0.349 ± 0.074 μm/min (567±120 dimers/min) with p (*Pf*, *Hs*) <0.0001, g=0.42. (**d**) *Pf* microtubules depolymerize at 6.62± 3.36 μm/min (10800±5460 dimers/min), *Hs* microtubules at 5.46±2.22 μm/min (8870±3610 dimers/min) with p (*Pf*, *Hs*) =0.0002, g=0.40. (**e**) Catastrophe frequencies are reported as the inverse of microtubule lifetimes. *Pf* microtubules catastrophe at 0.0047±0.0035 s^−1^, *Hs* microtubules at 0.0058±0.0012 s^−1^ with p (*Pf*, *Hs*) <0.0001, g=0.42. (**f**) Rescue frequencies are reported as the inverse of the duration of each depolymerization event. Events without a rescue are given a value of zero. *Pf* microtubules rescue at 0.0120±0.0251 s^−1^, *Hs* microtubules at 0.0494±0.0284 s^−1^, with p (*Pf*, *Hs*) <0.0001, g=1.40. Total time (T_total_) of observed microtubule growth and shrinkage and number of events (n) indicated.

### Microtubule-targeting compounds selectively inhibit *P. falciparum* but not human microtubule growth

Previous studies have proposed that inhibition of blood-stage *P. falciparum* cell growth by the herbicides oryzalin and amiprofos methyl (APM) (Fig. 5a) is the result of the inhibition of microtubule assembly, and that these compounds selectively inhibit parasite rather than host microtubules (Dempsey et al. 2013, Fennell et al. 2006, Lyons-Abbott et al. 2010). Indeed, compounds such as oryzalin and APM have been shown to selectively inhibit parasite growth in cell culture in comparison to mammalian cells (Fennell et al. 2006). However, these studies did not show a direct selective effect of the compounds on human versus *P. falciparum* microtubules. Therefore, we still lack mechanistic evidence that oryzalin and APM interfere with parasite microtubule dynamics, and more importantly that microtubule inhibition is parasite-specific. To test whether inhibition of microtubule growth is indeed the molecular mechanism of action and whether the compounds selectively interact with parasite over host microtubules, dynamic *P. falciparum* and human microtubules were assembled *in vitro* in the presence or absence of oryzalin and APM. Consistent with oryzalin being a microtubule destabilizing agent (Morejohn et al. 1987), oryzalin concentrations as low as 2.5 μM effectively inhibited *P. falciparum* microtubule growth. APM fully inhibited *P. falciparum* microtubule growth at 10 μM. Importantly, both conditions did not affect mammalian microtubule growth (Fig. 5b). Titrations of oryzalin showed that *P. falciparum* microtubule growth velocity decreased with increasing oryzalin concentration over the range of 0.5 – 2.5 μM oryzalin (Fig. 5c), whereas HEK293 microtubule polymerization remained unchanged at oryzalin concentrations up to at least 25 μM (Fig. 5d). At 6 μM tubulin, *Plasmodium* microtubule growth rate was reduced to 50% with an oryzalin concentration of only 2 μM, i. e. at substoichiometric concentrations (33% of the tubulin concentration). Thus, there is a considerable regime in which oryzalin selectively disrupts the microtubules of *Plasmodium* without affecting vertebrate microtubules. To test for the inhibition of blood-stage parasite growth, *P. falciparum* cultures were grown for 96 hours with serial dilutions of the two compounds. Average IC50 values for oryzalin and APM were 3.85±0.37 μM and 4.32±0.45 μM (Fig. 5e and f), respectively, which are close to previously published values (Fennell et al. 2006). Taken together, we have for the first time demonstrated that tubulin and microtubules purified from *P. falciparum* can be studied directly with immediate physiological relevance: oryzalin and APM both selectively inhibit *P. falciparum* over mammalian microtubule growth at substoichiometric concentrations *in vitro*. Given the proven pharmacological value of disrupting microtubule function (Florian and Mitchison, 2016), the availability of purified parasite tubulin together with the assays established here will allow for more systematic, mechanistic, and quantitative bottom-up screens for pathogen-specific microtubule inhibitors.

**Figure 5.**
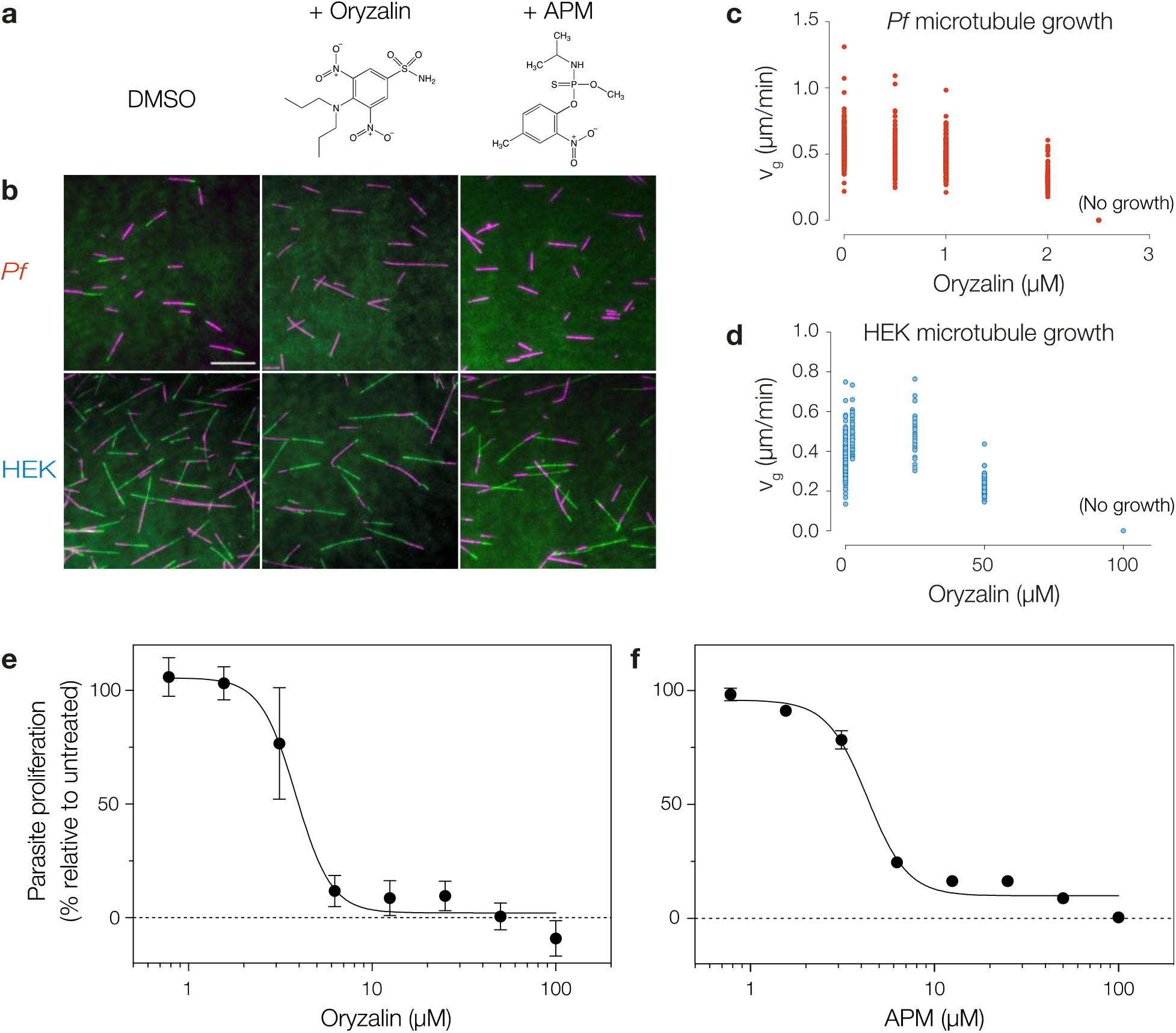
Oryzalin and APM are *P. falciparum*-specific microtubule inhibitors. (**a**) Structures of oryzalin and amiprofos methyl (APM) (**b**) Images of microtubules assembled from *P. falciparum* and HEK293 tubulin in 1 % DMSO, 2.5 μM oryzalin or 10 μM APM. All images were captured after 10 minutes of imaging at 37°C. Dynamic microtubules are shown in green, GMPCPP-stabilized microtubule seeds are shown in magenta. Scale bars: 10 μm. (**c**) *P. falciparum* and (d) HEK293 microtubule growth rates in the presence of increasing oryzalin concentrations with 6 μM tubulin. For *P. falciparum* microtubules, 192, 251, 144, and 134 growth events were measured at 0, 0.5, 1, and 2 μM oryzalin, respectively; no growth events were observed at or above 2.5 μM oryzalin. For HEK 293 microtubules, 223, 57, 33, and 62 growth events were measured at 0, 2.5, 25, and 50 μM oryzalin, respectively; no growth events were observed at 100 μM oryzalin. Representative dose-response curves of static *P. falciparum* cultures grown in the presence of oryzalin (**e**) and APM (**f**). Each point represents parasite proliferation relative to an untreated (100%) and chloroquine-treated (0%) control. Error bars indicate SD between replicates, and where not visible, are smaller than the symbol. Oryzalin titration data represent nine data points averaged from three independent experiments; APM titration data represent data averaged from three technical replicates.

## DISCUSSION

With the idea to selectively disrupt the microtubules of protozoan parasites without affecting vertebrate microtubules, we report the purification of *P. falciparum* tubulin from infected red blood cells, which is assembly-competent. The advantages of choosing tubulin as a target candidate are several-fold: (1) As tubulin is essential for all *Plasmodium* life-cycle stages, a drug interfering with microtubule function will thus be likely to target not only the asexual erythrocytic stage (Spreng et al. 2019) but also the invasion of erythrocytes by newly formed merozoites (Bell 1998). (2) Importantly, tubulin genes are not prone to mutations and thus resistance to microtubule inhibitors may be slow to develop. Although tubulin genes can mutate, this happens at a fitness cost in parasites (Ma et al. 2007). (3) Finally, there is a broad understanding of formulation compatibility and safety of microtubule-disruptive drugs, which can be built on (Florian and Mitchison, 2016).

While we here use a purification strategy that relies on the universal affinity of the yeast TOG-domain to tubulin (Widlund et al. 2011), we argue that the degree of amino acid conservation between human and *P. falciparum* α- and β-tubulin (83.7 and 88.5 % identity, respectively, Supplementary Fig. 3b) is sufficiently divergent to make *Plasmodium* tubulin an attractive target for drug discovery. Although tubulin is highly conserved in eukaryotes, single point mutations have been shown to be sufficient to increase sensitivity to microtubule-targeting agents (Kanakkanthara et al. 2015). Until now, six distinct ligand-binding sites on mammalian tubulin have been identified (as reviewed in Steinmetz and Prota 2018), most of them on β-tubulin (Supplementary Fig. 4). What about the oryzalin binding site? Computational and genetic resistance studies suggest that the oryzalin binding site is located on the α-tubulin subunit (Morrissette et al. 2004, Mitra and Sept 2006, Ma et al. 2007, Lyons-Abbott et al. 2010). However, when comparing mutation sites that are suggested to confer oryzalin resistance (Ma et al. 2007, Lyons-Abbott et al. 2010), we find many to be conserved in human α-tubulin (Supplementary Fig. 5). Clearly, further studies are required to evaluate the selective toxicity of oryzalin to Apicomplexan tubulin. The availability of purified functional parasite tubulin, however, will now allow us to determine the structure of *Plasmodium* tubulin, for example by using cryo-electron microscopy (cryo-EM), which will lay a solid ground for future structure-based drug design.

Microtubules assembled from *P. falciparum* tubulin, which we purified from parasite cells cultured in human erythrocytes, are made of only two isoforms (α1β), carry a monoglutamylation and show similar characteristics of dynamic instability as mammalian microtubules. While the reported isoform composition is consistent with previous antibody-based experiments (Delves et al. 1989, Holloway 1989, Holloway 1990, Rawlings 1992) and expression data (Toenhake et al. 2018, Josling et al. 2015), we do not find the purified tubulin to be polyglutamylated but a small fraction of β-tubulin to be monoglutamylated. This seems to be in contrast to recent immunofluorescence data, which show subpellicular microtubules to be polyglutamylated in *P. falciparum* schizonts (Bertiaux et al. 2020). One explanation for this discrepancy might be that subpellicular microtubules are not the predominant form of microtubules in schizonts and thus only represent a minor population in our purified tubulin. Indeed, the same study shows that hemispindles are not polyglutamylated in schizonts (Bertiaux et al. 2020).

While malaria remains one of the most devastating diseases, we hope that our findings encourage the use of parasite tubulins for the future identification of selective microtubule-binding drugs not only as an antimalarial but as potential cure for numerous human pathogens. In addition, recent estimates indicate that the plant kingdom comprises at least 500,000 species, of which less than 10 percent have been phytochemically investigated for pharmacological applications, leaving wide open the possibility that many new compounds that may target microtubules remain to be discovered (Zhang & Kanakadhara 2020).

## METHODS

### Preparation of GMPCPP-stabilized microtubule seeds

To serve as nucleation templates for *in vitro* microtubule assembly, stabilized microtubule seeds were prepared using GMPCPP, a slowly-hydrolyzing GTP analogue that prevents rapid microtubule depolymerization (Hyman et al. 1992). Additionally, biotinylated tubulin was incorporated to facilitate adhesion to the surface of neutravidin-functionalized coverslips, and labelled tubulin was used to visualize the seeds. A seed assembly reaction was prepared on ice by mixing bovine brain tubulin (final concentration 2.8 μM), fluorescently labelled tubulin (final concentration 0.4 μM), biotin-labeled tubulin (final concentration 0.8 μM), MgCl2 (final concentration 2 mM, and GMPCPP (final concentration 1 mM) in 1x BRB80 in a microreaction tube. The reaction was left on ice for 10 minutes to allow for nucleotide exchange, and then transferred to a ThermoMixer (Eppendorf) and incubated at 37°C for 2 hours. The reaction was then removed from the incubator and microtubules were gently resuspended by flicking the tube. Using a 200 μL pipette with cut-off tip to avoid shearing, the suspension was gently layered over 100 μL of a pre-warmed (37°C) 1x BRB80 solution supplemented with 60 % v/v glycerol and 0.5 mM GMPCPP in a 230 μL thick-walled centrifuge tube (Beckman-Coulter). Care was taken to maintain the two distinct liquid phases. The reaction was centrifuged in a pre-warmed (37°C) TLA-100 rotor at 22,0000 xg for 10 minutes at 35°C. The top layer was removed, and the cushion interface was washed with warm 1x BRB80, after which the cushion was removed, and the pellet was gently washed twice with warm 1x BRB80. Finally, the pellet was resuspended in warm 1x BRB80 supplemented with 1 mM DTT using a cut-off pipette tip, separated into 5 μL working aliquots, flash-frozen in liquid nitrogen and stored at −80°C.

### TIRF assays, image acquisition, and data processing

Flow chambers were constructed with glass coverslips passivated with trimethylchlorosilane (as described in Hirst et al. 2020b) and mounted onto passivated glass slides using thin strips of parafilm. Chambers were functionalized by perfusing 20 μL of 100 mg/mL Neutravidin in 1x BRB80 through the chamber and incubating for 5 min at room temperature. The chamber was rinsed twice with 20 μL 1x BRB80, twice with a blocking buffer consisting of 1 % w/v Pluronic F-127 (Sigma-Aldrich) in 1x BRB80, and incubated at room temperature for 15 min. Wash buffer containing 1 mg/mL Κ-casein in 1x BRB80 was flowed through the chamber followed by 2 x 20 μL of GMPCPP-stabilized microtubule seeds containing 10 % Atto565- or Atto647-labeled and 20 % biotin-labeled tubulin suspended in wash buffer.

Polymerization reactions were carried out at 37°C in 1x BRB80 buffer supplemented with 1 mg mL k-casein, 1 % β-mercaptoethanol, 2 mM Trolox, 2.5 mM PCA, 25 nM PCD, and 0.15 % methylcellulose at different concentrations of purified tubulin with 10 % Atto565- or Atto488-labeled porcine brain tubulin. In the case of 6 μM tubulin reactions, glycerol was added to a final concentration of 10% v/v to promote microtubule growth at low tubulin concentrations. 30 - 50 μL of reaction solution were perfused through the chamber, then both ends were sealed with silicone grease. The slide was mounted on the objective and left for 10 min to allow the temperature to equilibrate before imaging.

Images were taken on an inverted Nikon Eclipse Ti-E microscope with a motorized TIRF angle, a Nikon Plan Apochromat 100x/1.49NA oil immersion objective lens, and a Photometrics Prime 95B sCMOS camera. Atto488-labeled microtubules were imaged with a 488 nm laser, Atto565-labeled microtubules were imaged with a 561 nm laser and Atto647-labeled microtubule seeds were imaged with a 647 nm laser. Time-lapse images were taken at a frame rate of 0.2 fps with an exposure time of 100 - 200 ms. Recording was controlled with the Nikon ND Acquisition software.

Microtubule dynamics were measured by producing kymographs using the Multi Kymograph function of the FIJI image analysis software (Schindelin et al. 2012) and manually fitting lines to growth and shrinkage events according to (Hirst et al. 2020b). Growth and shrinkage velocities were calculated from the slopes of the fitted lines. Catastrophe frequencies were calculated as the inverse of the mean of microtubule lifetimes. Rescue frequencies were calculated as the inverse of the mean duration of depolymerization events; events without a rescue were assigned a value of 0.

### Plasmodium falciparum cell culture

The chloroquine (CQ)-sensitive 3D7 wild-type strain of *P. falciparum* was used for all cell culture and purification of *Pf* tubulin. Parasites were cultured in complete culture media (CCM) (RPMI supplemented with 3 mM L-glutamine, 25 mM HEPES, 20 mM glucose, 24 μg/mL gentamycin, 200 μM hypoxanthine, and 0.6 % AlbuMAX II bovine serum albumin (Gibco)). Routine cultures were maintained at 3 – 4 % haematocrit in a 37ºC incubator with a gas mixture of 1 % CO_2_, 3 % O_2_, and 96 % N_2_ on a shaking platform. Parasitaemia was monitored every 24 - 48 hours by Giemsa stain and cultures were split when the parasitaemia reached 1 – 10 %.

### Sorbitol synchronization of *P. falciparum* cultures

Cultures with >1 % ring-stage parasitaemia were centrifuged at 500 x g for 5 minutes at room temperature and the supernatant was discarded. Pellets were resuspended in 10 pellet volumes of pre-warmed (37°C) 5 % w/v sorbitol and incubated at 37°C for 15 minutes. The cell suspension was centrifuged again for 5 minutes at 500 x g at room temperature, the supernatant was discarded, and the pellet was resuspended in complete RPMI to the desired haematocrit to continue culturing.

### Purification of *P. falciparum* tubulin

To purify tubulin from *P. falciparum*, parasite cells were first isolated from host red blood cells by saponin lysis. Synchronized cultures at >5 % parasitaemia were pelleted at 500 x g at 4°C in a Beckmann 5810 R centrifuge and the supernatant was discarded. The pellet was resuspended on ice in an equal volume of saponin lysis buffer and incubated on ice for 5 minutes with occasional inversion of the tube. The pellet was centrifuged for 5 minutes at 500 x g at 4°C, the supernatant was discarded, and the pellet was resuspended in a volume of 1x PBS equivalent to the total original saponin lysis suspension. The pellet was again centrifuged and the 1x PBS wash was repeated twice. Finally, the pellet was resuspended in a minimal volume of 1x PBS (1 mL or less), flash frozen, and stored at −80°C. On the day of tubulin purification, aliquots were quickly thawed, pooled, resuspended in 1x BRB80, and supplemented with 100 μM GTP, 2 mM DTT, and 1x protease inhibitors. Cells were sonicated at full power at 0.5 second intervals totaling 30 seconds with a sonicator probe (Sonorex GM 2070) and then centrifuged for 10 minutes at 80,000 rpm at 4°C in an MLA-80 rotor. The supernatant was loaded at 0.5 CV/min onto a TOG-column preequilibrated with 1× BRB80. The flow rate was then changed to 1 CV/min for the following wash steps: 1) 10 CV of 1× BRB80, 100 μM GTP; 2) 3 CV of 1× BRB80, 100 μM GTP, 10 mM MgCl_2_, 5 mM ATP followed by a 15-min incubation; 3) 5 CV of 1× BRB80 and 100 μM GTP. The tubulin was eluted with 3 CV of 1× BRB80, 100 μM GTP, and 500 mM (NH_4_)_2_SO_4_. Pooled fractions were desalted into 1× BRB80 and 10 μM GTP with a PD10 desalting column and concentrated using an Amicon Ultra 30K MWCO centrifugal filter (Millipore). Tubulin was aliquoted and snap frozen in liquid nitrogen.

### Purification of HEK tubulin

HEK293 cells were pelleted and resuspended in an equal volume of 1× BRB80 containing 3 U of benzonase, 1 mM DTT, and protease inhibitors. Cells were lysed by douncing on ice and the lysate was cleared by centrifugation at 80,000 rpm (440,000 x g) in a pre-cooled MLA-80 rotor at 2°C for 30 minutes and then filtered through a 0.45-μm Milliex-HV polyvinylidene fluoride membrane (Millipore, Bedford, MA). The filtrate was loaded onto an equilibrated TOG-column. Tubulin was eluted, desalted, concentrated and flash-frozen in liquid nitrogen as described above.

### Purification of bovine brain tubulin

Frozen bovine brain tissue stored at −80°C was pulverized using a mortar and pestle on dry ice and then suspended in BRB80 supplemented with benzonase, 1 mM DTT, protease inhibitors, and 10 μg/mL Cytochalasin D at a ratio of 1 mL buffer: 1 g brain tissue on ice. The tissue suspension was sonicated at full power at 0.5 second intervals totalling one minute with a sonicator probe (Sonorex GM 2070) and then further homogenized using 20 strokes with a cell douncer on ice. The lysate was centrifuged for 10 minutes at 80,000 rpm at 4°C in an MLA-80 rotor, the supernatant was loaded onto an equilibrated TOG affinity column, and the tubulin purification proceeded as previously described.

### Dose-response assays

Synchronous blood-stage shaking cultures between 1 % and 7 % parasitaemia were used for dose-response assays. A 10 mL aliquot of culture suspension was removed from the culture flask and the iRBCs were washed by centrifuging at 500 x g for five minutes, removing the supernatant, resuspending the pellet in CCM, and incubating for 1 hour in a 37°C water bath. In parallel, 2 mL of fresh uninfected RBCs (uRBCs) were washed by adding the cells to 10 mL CCM and incubating at 37°C for 1 hour. Tubes containing cell suspensions were inverted every 15 – 20 min to keep the cells suspended. While the cells were incubating during the wash, medium containing the inhibitor of interest was prepared by diluting the inhibitor stock in CCM and sterile filtering with a syringe attached to a 0.2 μM syringe filter. The working solution inhibitor concentration was 2x the highest concentration tested during the assay. The volume of CCM used corresponded to the number of rows of the 96-well culture plate used for each inhibitor. Generally, 1.6 mL of medium was prepared each time, corresponding to 200 μL per row for 3 technical replicates plus 1 mL extra in case any solution was lost during filtering. Additionally, CCM containing 1 mM chloroquine (CQ) was prepared as a positive control in the same manner to a final volume of 1 mL per plate. Dose-response assays were carried out in 96-well culture plates with the rows labelled A – H and the columns labelled 1 – 12. Only rows B – G and columns 2 – 11 contained samples; all peripheral wells contained 200 μL RPMI to prevent evaporation from experimentally important wells. To prepare the wells for cell culture, 100 μL of CQ working solution were added to the remaining wells of column 2, 200 μL of each inhibitor working solution were added to column 4, and 100 μL of CCM were added to the remaining wells. The inhibitor solution of column 4 was then serially diluted at a ratio of 1:2 from column 4 to column 11, leaving a volume of 100 μL in each well. To prepare the working iRBC cell suspension, the iRBC and uRBC suspensions were removed from the water bath, centrifuged at RT for 5 min at 500 xg, and the supernatant was discarded. iRBCs and uRBCs were resuspended in CCM at an appropriate ratio to achieve 2 % hematocrit at 0.5 % parasitaemia. 100 μL of the cell suspension were added to each well to achieve a final culture volume of 200 μL at 1 % hematocrit. Finally, plates were incubated in sealed humidified chambers with 3 % oxygen and 1 % CO2 and incubated at 37°C without shaking for 96 h. Parasite growth was quantified by measuring the quantity of DNA using the DNA dye SYBR Safe. At the end of 96 hours, cell cultures were resuspended in their wells transferred to a new 96-well plate, and mixed with a lysis buffer (20 mM TRIS, 5 mM EDTA, 0.008 % w/v saponin, 0.08 % v/v Triton X-100, pH 7.5) containing 0.02 % v/v SYBR Safe dye. Fluorescence at the 490 nm excitation and 520 nm emission wavelengths was measured using a FLUOstar OPTIMA plate reader (BMG Labtech). The signals were normalized to a scale of 0 – 100 % based on the mean signals of the CQ-treated (0 %) and untreated (100 %) wells. To determine IC50 values, the data were fit to a four parameters logistic regression model using the GraphPad Prism software.

### Intact protein mass spectrometry

The purified *P. falciparum* tubulin was analyzed using the Ultimate 3000 liquid chromatography system connected to a Q Exactive HF mass spectrometer via the ion max source with HESI-II probe (Thermo Scientific). The following MS source parameters were used: spray voltage 3.6 kV, capillary temperature 320°C, sheath gas 10, auxiliary gas 4, S-lens RF level 60, intact protein mode on. For the analysis 2 μL (approx. 1.3 μg of total protein) of the sample were desalted and concentrated by injection on a reversed-phase cartridge (MSPac DS-10, 2.1×10 mm, Thermo Scientific) at 60°C using buffer A (0.1 % formic acid, 5 % acetonitrile in water) at a constant flow rate of 22 μL/min for 3 min. This was followed by a short linear gradient of 5 % – 95 % buffer B (0.1 % formic acid in 80 % acetonitrile, 20 % water) within 10 min followed by washing and re-equilibration. Full MS spectra were acquired using the following parameters: mass range m/z 600–2500, resolution 15,000, AGC target 3×10^6^, μscans 5, maximum injection time 200 ms. MS raw data were analysed using BioPharma Finder (version 3.2, Thermo Scientific). First, an averaged spectrum over the chromatographic peak was generated followed by spectral deconvolution using a deconvolution mass tolerance of 5 ppm and a relative abundance threshold of 10 %.

### Peptide mass fingerprint analysis by MALDI-MS

Purified *Plasmodium* tubulin was separated by SDS-PAGE (as exemplified in Fig. 1b) and bands of α- and ß-tubulin were excised separately and subjected to trypsin in-gel digestion as described previously (Shevchenko et al. 1996). Peptide masses were recorded by matrix-assisted laser desorption ionization-time of flight mass spectrometry (MALDI-TOF-MS) using an Ultraflex-II TOF/TOF instrument (Bruker Daltonics, Bremen, Germany) equipped with a 200 Hz solid-state Smart beam™ laser. We used -cyano-4-hydroxycinnamic acid (CHCA) as the matrix and applied the protein digest samples using the dried droplet technique. The mass spectrometer was operated in the positive reflector mode in the m/z range of 6004,000. Database searches were performed using Mascot (Matrix Science Ltd., http://www.matrixscience.com), with mass tolerance typically set at 75 ppm and one missed cleavage allowed. MS/MS spectra of selected peptides were acquired using the LIFT mode (Suckau et al. 2003).

### Antibodies

A monoclonal anti-α-tubulin (SIGMA, T5168, Clone B-5-1-2) was used in western blots as a loading control. In addition, we used sheep polyclonal ATN02 also directed against α-tubulin and β-tubulin (Sigma T7816). K40: Acetylated lysine 40 of α-tubulin (Sigma Aldrich T7451 clone 6-11 B-1). Poly-Glu: Polyglutamylation of acidic residues (Adipogen gt335). Tyr: Tyrosinated C terminus of tubulin (Abcam ab6160). Detyr: Tubulin C terminus with tyrosine removed (Abcam ab48389). HSP70: *P. falciparum* HSP70 (LSBio LS-C109068). Secondary antibodies were HRP-conjugated anti-rabbit IgG (Proteintech 0001-2) and anti-mouse IgG (Proteintech 0001-1) from goat.

### Compounds

Stock solutions of 100 mM oryzalin and 100 mM APM were prepared in 100 % dimethyl sulfoxide (DMSO), filter sterilized, and stored in single-use aliquots at −20°. On the day of a TIRF experiment, a single aliquot was further diluted in DMSO such that addition of compound to the master mix resulted in a final DMSO concentration of no more than 1% v/v.

### Statistical analysis

In each figure legend, details about the quantifications have been provided, including the number of events measured (n), the mean/median values, the SD/SEM. In addition, information about the statistical tests used for measuring significance and interpretation of p values is provided. P values greater than 0.05 are represented by “ns”. A single * indicates a p value ≤ 0.05, ** indicates p values ≤ 0.01, *** indicates p values ≤ 0.001, and **** indicates p values ≤ 0.0001. In addition to the significance, we also indicate the effect size by calculating Hedges’ g. An effect size between 0.20 – 0.50 was considered small, while effect sizes between 0.51 – 0.80 were considered medium and g > 0.81 was considered large. For statistical analysis and plotting in this paper, we utilized Graphpad Prism version 8.0 for Mac OS X, GraphPad Software, La Jolla California USA, https://www.graphpad.com.

## Supporting information

Hirst_Suppl

## ACKNOWLEDGEMENTS

The authors thank all past and current members of the Reber lab, in particular Sebastian Reusch and Soma Zsoter for purifying HEK293 tubulin. We thank the AMBIO imaging facility (Charite, Berlin) for imaging support. For mass spectrometry, we would like to acknowledge the assistance of the Core Facility BioSupraMol supported by the Deutsche Forschungsgemeinschaft (DFG). We are grateful to the Baum and Matuschewski labs for discussions, helpful advice and providing essential materials. We further thank Ylva Veith for maintaining *P. falciparum* cultures. We would like to thank the Canberra branch of the Australian Red Cross Lifeblood for the provision of red blood cells. We thank Stefan Florian for critical comments on the manuscript. This work was supported by the Alliance Berlin Canberra ‘Crossing Boundaries: Molecular Interactions in Malaria’, which is co-funded by a grant from the Deutsche Forschungsgemeinschaft (DFG) for the International Research Training Group (IRTG) 2290 and the Australian National University. S.R. further acknowledges funding by the IRI Life Sciences (Humboldt-Universität zu Berlin, Excellence Initiative/DFG).

## AUTHOR CONTRIBUTIONS

W.G.H. and S.R. conceived the project. W.G.H. performed all experiments and analysed the data. D.F. contributed purified tubulin. B.K. and C.W. performed and interpreted the mass-spectrometric analyses. Experiments with *P. falciparum* cultures were carried out in the lab of K.S. S.R. wrote the manuscript with input from all authors.

## ADDITIONAL INFORMATION

– Supplementary movie S1
– Figure S1-S5.

