## Supplementary material for "Purification of functional *Plasmodium falciparum* tubulin allows for the identification of parasite-specific microtubule inhibitors": Hirst_Suppl

- Supplementary Movie S1
- Supplementary Figures S1-S5

**Supplementary Movie S1:** TIRF microscopy time lapse (25 fps) of dynamic microtubules assembled from purified *P. falciparum* tubulin (green) grown from stabilized seeds (magenta) at 37°C and 6 μM tubulin for 20 minutes. Scale bar: 5 μm. Time from beginning of image capture is shown as min:sec. The movie corresponds to the still image shown in Figure 1e.

**Hirst et al. - Figure S1**

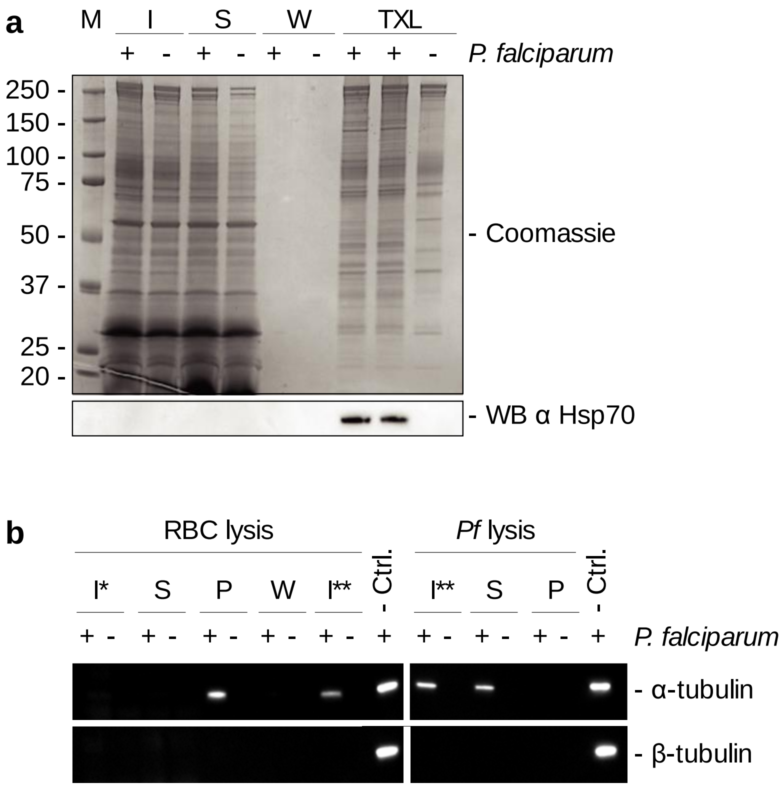


**Supplementary Figure 1:** Tubulin purification from *P. falciparum* infected red blood cells. (**a**) Sequential lysis of infected (+) and uninfected (-) erythrocytes followed by lysis of *P. falciparum* cells. M: Molecular weight marker (kDa), I: Saponin lysis input, S: Supernatant, W: Wash, TXL: Triton-X lysis. Lower panel: As an additional control, the host cell lysis was monitored by western blot with a *P. falciparum*-specific Hsp70 antibody to confirm that parasite cells remained intact until the final lysis step. (**b**) The presence of tubulin during sequential lysis of infected (+) and uninfected (-) erythrocytes was monitored using a pan-specific α-tubulin and a vertebrate-specific β-tubulin antibody. I*: Saponin lysis input, S: Supernatant, P: Pellet, W: PBS wash supernatant, I** *P. falciparum* lysis input, Ctrl: HeLa tubulin control.

### Hirst et al. - Figure S2

**a**

```
α1|Q6ZLZ9|          MREVISIHVGQAGIQVGNACWELFCLEHGIQPDGQMPSDKASRANDDAFNTFFSETGAGK 060
α2|Q8IFP3|          MREVISIHVGQAGIQIGNACWELFCLEHGIQPDGQMPSDQVVAGGDDAFNTFFSETGAGK 060
β |Q7KQL5|          MREIVHIQAGQCGNQIGAKFWEVISDEHGIDPSGTYCGDSDL--QLERVDVFYNEATGGR 058
                    ***:: *:.**.* *:*   **::. ****:*.*   .*.      : .:.*:.*: .*:

α1|Q6ZLZ9|          HVPRCVFVDLEPTVVDEVRTGTYRQLFHPEQLISGKEDAANNFARGHYTIGKEVIDVCLD 120
α2|Q8IFP3|          HVPRCVFVDLEPTVVDEVRTGTYRQLFHPEQLISGKEDAANNFARGHYTIGKEIVDVCLD 120
β |Q7KQL5|          YVPRAILMDLEPGTMDSVRAGPFGQLFRPDNFVFGQTGAGNNWAKGHYTEGAELIDAVLD 118
                    :***.:::**** .:*.**:* : ***:*:::: *: .*.**:*:**** * *::*. **

α1|Q6ZLZ9|          RIRKLADNCTGLQGFLMFSAVGGGTGSGFGCLMLERLSVDYGKKSKLNFCCWPSPQVSTA 180
α2|Q8IFP3|          RVRKLADNCTGLQGFLMFNAVGGGTGSGLGCLLLERLAIDYGKKSKLNFCSWPSPQVSTA 180
β |Q7KQL5|          VVRKEAEGCDCLQGFQITHSLGGGTGSGMGTLLISKIREEYPDRIMETFSVFPSPKVSDT 178
                     :** *:.*  **** :  ::*******:* *::.::  :* .:   .*. :***:** :

α1|Q6ZLZ9|          VVEPYNSVLSTHSLLEHTDVAIMLDNEAIYDICRRNLDIERPTYTNLNRLIAQVISSLTA 240
α2|Q8IFP3|          VVEPYNSVLSTHSLLEHTDVAIMLDNEAIYDICKKNLDIERPTYTNLNRLIAQVISSLTA 240
β |Q7KQL5|          VVEPYNATLSVHQLVENADEVQVIDNEALYDICFRTLKLTTPTYGDLNHLVSAAMSGVTC 238
                    ******:.**.*.*:*::* . ::****:**** :.*.:  *** :**:*:: .:*.:*.

α1|Q6ZLZ9|          SLRFDGALNVDVTEFQTNLVPYPRIHFMLSSYAPVVSAEKAYHEQLSVSEITNSAFEPAN 300
α2|Q8IFP3|          SLRFDGALNVDVTEFQTNLVPYPRIHFMLSSYAPIISAEKAYHEQLSVSEITNSAFEPAS 300
β |Q7KQL5|          SLRFPGQLNSDLRKLAVNLIPFPRLHFFMIGFAPLTSRGSQQYRALTVPELTQQMFDAKN 298
                    **** * ** *: :: .**:*:**:**:: .:**: *  .  :. *:* *:*:. *:  .

α1|Q6ZLZ9|          MMAKCDPRHGKYMACCLMYRGDVVPKDVNAAVATIKTKRTIQFVDWCPTGFKCGINYQPP 360
α2|Q8IFP3|          MMAKCDPRHGKYMACCLMYRGDVVPKDVNAAVATIKTKRSIQFVDWCPTGFKCGINYQPP 360
β |Q7KQL5|          MMCASDPRHGRYLTACAMFRGRMSTKEVDEQMLNVQNKNSSYFVEWIPHNTKSSVCDIPP 358
                    **. .*****:*::.* *:** :  *:*:  : .::.*.:  **:* * . *..:   **

α1|Q6ZLZ9|          TVVPGGDLAKVMRAVCMISNSTAIAEVFSRMDQKFDLMYAKRAFVHWYVGEGMEEGEFSE 420
α2|Q8IFP3|          TVVPGGDLAKVMRAVCMISNSTAIAEVFSRMDQKFDLMYAKRAFVHWYVGEGMEEGEFSE 420
β |Q7KQL5|          KG--------LKMAVTFVGNSTAIQEMFKRVSDQFTAMFRRKAFLHWYTGEGMDEMEFTE 410
                    .         :  ** ::.***** *:*.*:.::*  *: ::**:***.****:* **:*

α1|Q6ZLZ9|          AREDLAALEKDYEEVGIESNEAEGEDEGYEADY-- 453
α2|Q8IFP3|          AREDLAALEKDYEEVGIESNDGEGEDEGYE----- 450
β |Q7KQL5|          AESNMNDLVSEYQQYQDATAEEEGEFEEEEGDVEA 445
                    *..::  * .:*::    : : *** *  *
```

**b**

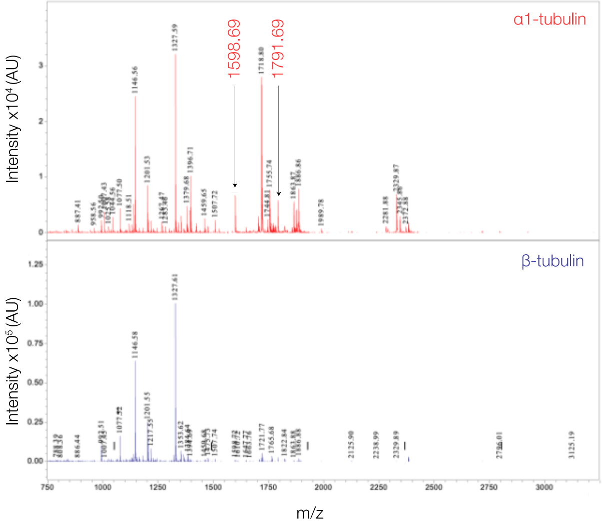


**c**

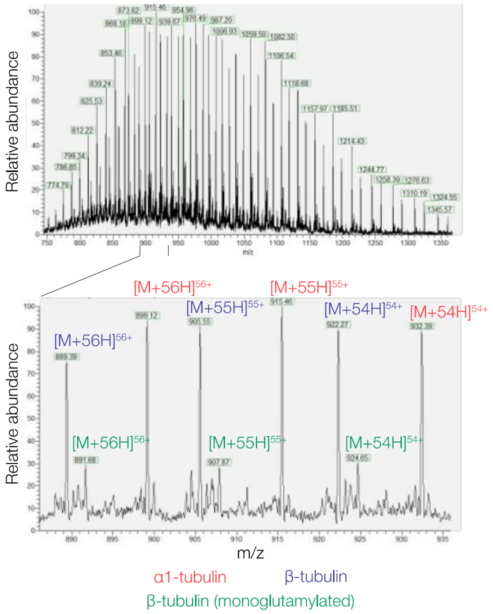

**Supplementary Figure 2**: *P. falciparum* tubulin isoforms. (**a**) Multiple sequence alignment of all three *P. falciparum* tubulins aligned with the T-Coffee algorithm. Peptides unique to the α1-isoform that were identified by mass spectrometry are highlighted in red. (**b**) Peptide mass fingerprint analysis by MALDI-MS. *P. falciparum* α1-tubulin peptide masses are indicated in red and β-tubulin masses are in blue. The α1-tubulin mass fingerprint from the trypsin digest shows peaks corresponding to peptides that are shared by both α1- and α2-tubulins with the exception of the peaks at 1,791.7 Da, which is a unique peptide corresponding to amino acids 44–60 of α1-tubulin and 1,598.7 corresponding to the tryptic peptide 340–352 of α1-tubulin. A Thr→Ser mutation is found in α2-tubulin in this region of the sequence and there was no signal at the corresponding mass of 1,584.7 Da. (**c**) Mass spectrum of purified *P. falciparum* tubulin recorded under denaturing conditions. The whole mass spectrum (top) shows a broad charge state distribution. The zoomed-in view (bottom) shows a representative fraction of highly abundant charge states (54+ to 56+) indicating the presence of at least three different protein species. Signals matching to α1-tubulin, β-tubulin and monoglutamylated β-tubulin are highlighted in the legend. The deconvoluted spectrum is shown in Figure 2A of the main text.

**Hirst et al. - Figure S3**

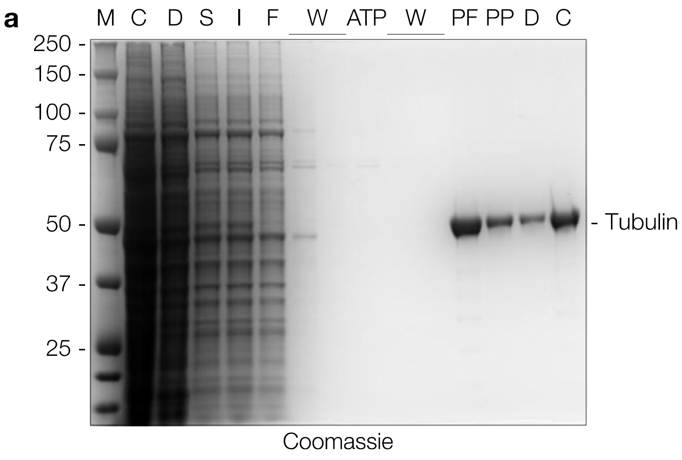


**b** % of pairwise protein identity

| Human | *Pf* α1 | *Pf* α2 | *Pf* β |
|---|---|---|---|
| Q71U36 (Tubulin α-1A chain) | 83.7 | 82.9 | 41.2 |
| P68363 (Tubulin α-1B chain) | 83.4 | 83.1 | 41.6 |
| Q9BQE3 (Tubulin α-1C chain) | 83.2 | 82.4 | 41.0 |
| P0DPH7 (Tubulin α-3C chain) | 83.3 | 82.9 | 40.3 |
| P0DPH8 (Tubulin α-3D chain) | 83.3 | 82.9 | 40.3 |
| Q6PEY2 (Tubulin α-3E chain) | 82.2 | 81.6 | 40.2 |
| P68366 (Tubulin α-4A chain) | 81.7 | 81.3 | 40.5 |
| Q9NY65 (Tubulin α-8 chain) | 78.6 | 78.4 | 40.3 |
| P07437 (Tubulin β-5 chain) | 41.1 | 41.3 | 88.5 |
| Q9H4B7 (Tubulin β-1 chain) | 38.9 | 38.1 | 73.8 |
| Q13885 (Tubulin β-2A chain) | 40.3 | 40.8 | 88.1 |
| Q9BVA1 (Tubulin β-2B chain) | 40.3 | 40.8 | 88.3 |
| Q13509 (Tubulin β-3 chain) | 40.3 | 40.3 | 84.9 |
| P04350 (Tubulin β-4A chain) | 40.6 | 40.7 | 88.1 |
| P68371 (Tubulin β-4B chain) | 40.8 | 40.8 | 88.5 |
| Q9BUF5 (Tubulin β-6 chain) | 40.2 | 41.1 | 83.9 |
| Q3ZCM7 (Tubulin β-8 chain) | 39.1 | 39.7 | 82.7 |

**Supplementary Figure 3**: Purification of human tubulin from HEK293 cells. (**a**) Coomassie-stained SDS-PAGE of the individual purification steps from HEK293 lysate. M: Marker, C: HEK293 cells, D: dounced HEK293 cells, S: Supernatant, I: Input, F: Flowthrough, W: Wash, ATP: ATP wash, PF: Peak fraction, PP: Pooled fractions, D: Desalt, C: Concentrated. (**b**) Pairwise protein identity between *P. falciparum* α1- (Uniprot ID Q6ZLZ9), α2- (Uniprot ID Q8IFP3), and β-tubulin (Uniprot ID Q7KQL5) and human tubulin isoforms as calculated by the Needleman-Wunsch algorithm.

### Hirst et al. - Figure S4

```
                                                1        10        20        30        40        50
P. falciparum Q7KQL5 (Tubulin β chain)          MREIVHIQAGQCGNQIGAKFWEVISDEHGIDPSGTYCGDSDLQLERVDVFYNEATGGRY
Human Q13885  (Tubulin β-2A chain)              MREIVHIQAGQCGNQIGAKFWEVISDEHGIDPTGSYHGDSDLQLERINVYYNEAAGNKY
Human P04350  (Tubulin β-4A chain)              MREIVHLQAGQCGNQIGAKFWEVISDEHGIDPTGTYHGDSDLQLERINVYYNEATGGNY
Human P68371  (Tubulin β-4B chain)              MREIVHLQAGQCGNQIGAKFWEVISDEHGIDPTGTYHGDSDLQLERINVYYNEATGGKY
Human Q9BVA1  (Tubulin β-2B chain)              MREIVHIQAGQCGNQIGAKFWEVISDEHGIDPTGSYHGDSDLQLERINVYYNEATGNKY
Human Q13509  (Tubulin β-3 chain)               MREIVHIQAGQCGNQIGAKFWEVISDEHGIDPSGNYVGDSDLQLERISVYYNEASSHKY

        60        70        80        90        100       110       120       130
Q7KQL5  VPRAILMDLEPGTMDSVRAGPFGQLFRPDNFVFGQTGAGNNWAKGHYTEGAELIDAVLDVVRKEAEGCDCLQGFQITHSI
Q13885  VPRAILVDLEPGTMDSVRSGPFGQIFRPDNFVFGQSGAGNNWAKGHYTEGAELVDSVLDVVRKESESCDCLQGFQLTHSI
P04350  VPRAVLVDLEPGTMDSVRSGPFGQIFRPDNFVFGQSGAGNNWAKGHYTEGAELVDAVLDVVRKEAESCDCLQGFQLTHSI
P68371  VPRAVLVDLEPGTMDSVRSGPFGQIFRPDNFVFGQSGAGNNWAKGHYTEGAELVDSVLDVVRKEAESCDCLQGFQLTHSI
Q9BVA1  VPRAILVDLEPGTMDSVRSGPFGQIFRPDNFVFGQSGAGNNWAKGHYTEGAELVDSVLDVVRKESESCDCLQGFQLTHSI
Q13509  VPRAILVDLEPGTMDSVRSGAFGHLFRPDNFIFGQSGAGNNWAKGHYTEGAELVDSVLDVVRKECENCDCLQGFQLTHSI

        140       150       160       170       180       190       200       210
Q7KQL5  GGGTGSGMGTLLISKIREEYPDRIMETFSVFPSPKVSDTVVEPYNATLSVHQLVENADEVQVIDNEALYDICFRTLKLTT
Q13885  GGGTGSGMGTLLISKIREEYPDRIMNTFSVMPSPKVSDTVVEPYNATLSVHQLVENTDETYSIDNEALYDICFRTLKLTT
P04350  GGGTGSGMGTLLISKIREEFPDRIMNTFSVVPSPKVSDTVVEPYNATLSVHQLVENTDETYCIDNEALYDICFRTLKLTT
P68371  GGGTGSGMGTLLISKIREEYPDRIMNTFSVVPSPKVSDTVVEPYNATLSVHQLVENTDETYCIDNEALYDICFRTLKLTT
Q9BVA1  GGGTGSGMGTLLISKIREEYPDRIMNTFSVMPSPKVSDTVVEPYNATLSVHQLVENTDETYCIDNEALYDICFRTLKLTT
Q13509  GGGTGSGMGTLLISKVREEYPDRIMNTFSVVPSPKVSDTVVEPYNATLSIHQLVENTDETYCIDNEALYDICFRTLKLAT

        220       230       240       250       260       270       280       290
Q7KQL5  PTYGDLNHLVSAAMSGVTCSLRFPGQLNSDLRKLAVNLIPFPRLHFFMIGFAPLTSRGSQQYRALTVPELTQQMFDAKNM
Q13885  PTYGDLNHLVSATMSGVTTCLRFPGQLNADLRKLAVNMVPFPRLHFFMPGFAPLTSRGSQQYRALTVPELTQQMFDSKNM
P04350  PTYGDLNHLVSATMSGVTTCLRFPGQLNADLRKLAVNMVPFPRLHFFMPGFAPLTSRGSQQYRALTVPELTQQMFDAKNM
P68371  PTYGDLNHLVSATMSGVTTCLRFPGQLNADLRKLAVNMVPFPRLHFFMPGFAPLTSRGSQQYRALTVPELTQQMFDAKNM
Q9BVA1  PTYGDLNHLVSATMSGVTTCLRFPGQLNADLRKLAVNMVPFPRLHFFMPGFAPLTSRGSQQYRALTVPELTQQMFDSKNM
Q13509  PTYGDLNHLVSATMSGVTTSLRFPGQLNADLRKLAVNMVPFPRLHFFMPGFAPLTARGSQQYRALTVPELTQQMFDAKNM

        300       310       320       330       340       350       360       370
Q7KQL5  MCASDPRHGRYLTACAMFRGRMSTKEVDEQMLNVQNKNSSYFVEWIPHNTKSSVCDIPPKGLKMAVTFVGNSTAIQEMFK
Q13885  MAACDPRHGRYLTVAAIFRGRMSMKEVDEQMLNVQNKNSSYFVEWIPNNVKTAVCDIPPRGLKMSATFIGNSTAIQELFK
P04350  MAACDPRHGRYLTVAAVFRGRMSMKEVDEQMLSVQSKNSSYFVEWIPNNVKTAVCDIPPRGLKMAATFIGNSTAIQELFK
P68371  MAACDPRHGRYLTVAAVFRGRMSMKEVDEQMLNVQNKNSSYFVEWIPNNVKTAVCDIPPRGLKMSATFIGNSTAIQELFK
Q9BVA1  MAACDPRHGRYLTVAAIFRGRMSMKEVDEQMLNVQNKNSSYFVEWIPNNVKTAVCDIPPRGLKMSATFIGNSTAIQELFK
Q13509  MAACDPRHGRYLTVATVFRGRMSMKEVDEQMLAIQSKNSSYFVEWIPNNVKVAVCDIPPRGLKMSSTFIGNSTAIQELFK

        380       390       400       410       420       430       440
Q7KQL5  RVSDQFTAMFRRKAFLHWYTGEGMDEMEFTEAESNMNDLVSEYQQYQDATAEEEGEFEEE-EGDVEA----
Q13885  RISEQFTAMFRRKAFLHWYTGEGMDEMEFTEAESNMNDLVSEYQQYQDATADEQGEFEEE-EGEDEA----
P04350  RISEQFTAMFRRKAFLHWYTGEGMDEMEFTEAESNMNDLVSEYQQYQDATAE-EGEFEEE-AEEEVA----
P68371  RISEQFTAMFRRKAFLHWYTGEGMDEMEFTEAESNMNDLVSEYQQYQDATAEEEGEFEEE-AEEEVA----
Q9BVA1  RISEQFTAMFRRKAFLHWYTGEGMDEMEFTEAESNMNDLVSEYQQYQDATADEQGEFEEE-EGEDEA----
Q13509  RISEQFTAMFRRKAFLHWYTGEGMDEMEFTEAESNMNDLVSEYQQYQDATAEEEGEMYEDDEEESEAQGPK
```

**Binding Sites** Taxol, Colchicine, Vinblastine, Mayatan, Laulilamide

**Supplementary Figure 4**: Multiple Sequence Alignment of β-tubulins including known drug binding sites. Multiple sequence alignment (MSA) of *P. falciparum* and human β-tubulin isoforms. Alignment was calculated with T-Coffee. Selected drug binding sites are highlighted: Taxol (orange), Colchicine (green), Vinblastine (purple), Mayatan (grey), and Laulimalide (pink).

### Hirst et al. - Figure S5

```
                                                  1          10        20        30        40        50
P. falciparum Q6ZLZ9 (Tubulin α-1 chain)          MREVISIHVGQAGIQVGNACWELFCLEHGIQPDGQMPSDKASRANDDAFNTFFSETGAG
P. falciparum Q8IFP3 (Tubulin α-2 chain)          MREVISIHVGQAGIQIGNACWELFCLEHGIQPDGQMPSDQVVAGGDDAFNTFFSETGAG
Human Q71U36 (Tubulin α-1A chain)                 MRECISIHVGQAGVQIGNACWELYCLEHGIQPDGQMPSDKTIGGGDDSFNTFFSETGAG
Human P68363 (Tubulin α-1B chain)                 MRECISIHVGQAGVQIGNACWELYCLEHGIQPDGQMPSDKTIGGGDDSFNTFFSETGAG
Human P0DPH7 (Tubulin α-3C chain)                 MRECISIHVGQAGVQIGNACWELYCLEHGIQPDGQMPSDKTIGGGDDSFNTFFSETGAG
Human P0DPH8 (Tubulin α-3D chain)                 MRECISIHVGQAGVQIGNACWELYCLEHGIQPDGQMPSDKTIGGGDDSFNTFFSETGAG
Human P68366 (Tubulin α-4A chain)                 MRECISVHVGQAGVQMGNACWELYCLEHGIQPDGQMPSDKTIGGGDDSFTTFFCETGAG

        60        70        80        90        100       110       120       130
Q6ZLZ9  KHVPRCVFVDLEPTVVDEVRTGTYRQLFHPEQLISGKEDAANNFARGHYTIGKEVIDVCLDRIRKLADNCTGLQGFLMFS
Q8IFP3  KHVPRCVFVDLEPTVVDEVRTGTYRQLFHPEQLISGKEDAANNFARGHYTIGKEIVDVCLDRVRKLADNCTGLQGFLMFN
Q71U36  KHVPRAVFVDLEPTVIDEVRTGTYRQLFHPEQLITGKEDAANNYARGHYTIGKEIIDLVLDRIRKLADQCTGLQGFLVFH
P68363  KHVPRAVFVDLEPTVIDEVRTGTYRQLFHPEQLITGKEDAANNYARGHYTIGKEIIDLVLDRIRKLADQCTGLQGFLVFH
P0DPH7  KHVPRAVFVDLEPTVVDEVRTGTYRQLFHPEQLITGKEDAANNYARGHYTIGKEIVDLVLDRIRKLADLCTGLQGFLIFH
P0DPH8  KHVPRAVFVDLEPTVVDEVRTGTYRQLFHPEQLITGKEDAANNYARGHYTIGKEIVDLVLDRIRKLADLCTGLQGFLIFH
P68366  KHVPRAVFVDLEPTVIDEIRNGPYRQLFHPEQLITGKEDAANNYARGHYTIGKEIIDPVLDRIRKLSDQCTGLQGFLVFH

        140       150       160       170       180       190       200       210
Q6ZLZ9  AVGGGTGSGFGCLMLERLSVDYGKKSKLNFCCWPSPQVSTAVVEPYNSVLSTHSLLEHTDVAIMLDNEAIYDICRRNLDI
Q8IFP3  AVGGGTGSGLGCLLLERLAIDYGKKSKLNFCSWPSPQVSTAVVEPYNSVLSTHSLLEHTDVAIMLDNEAIYDICKKNLDI
Q71U36  SFGGGTGSGFTSLLMERLSVDYGKKSKLEFSIYPAPQVSTAVVEPYNSILTTHTTLEHSDCAFMVDNEAIYDICRRNLDI
P68363  SFGGGTGSGFTSLLMERLSVDYGKKSKLEFSIYPAPQVSTAVVEPYNSILTTHTTLEHSDCAFMVDNEAIYDICRRNLDI
P0DPH7  SFGGGTGSGFASLLMERLSVDYGKKSKLEFAIYPAPQVSTAVVEPYNSILTTHTTLEHSDCAFMVDNEAIYDICRRNLDI
P0DPH8  SFGGGTGSGFASLLMERLSVDYGKKSKLEFAIYPAPQVSTAVVEPYNSILTTHTTLEHSDCAFMVDNEAIYDICRRNLDI
P68366  SFGGGTGSGFTSLLMERLSVDYGKKSKLEFSIYPAPQVSTAVVEPYNSILTTHTTLEHSDCAFMVDNEAIYDICRRNLDI

        220       230       240       250       260       270       280       290
Q6ZLZ9  ERPTYTNLNRLIAQVISSLTASLRFDGALNVDVTEFQTNLVPYPRIHFMLSSYAPVVSAEKAYHEQLSVSEITNSAFEPA
Q8IFP3  ERPTYTNLNRLIAQVISSLTASLRFDGALNVDVTEFQTNLVPYPRIHFMLSSYAPIISAEKAYHEQLSVSEITNSAFEPA
Q71U36  ERPTYTNLNRLIGQIVSSITASLRFDGALNVDLTEFQTNLVPYPRIHFPLATYAPVISAEKAYHEQLSVAEITNACFEPA
P68363  ERPTYTNLNRLISQIVSSITASLRFDGALNVDLTEFQTNLVPYPRIHFPLATYAPVISAEKAYHEQLSVAEITNACFEPA
P0DPH7  ERPTYTNLNRLIGQIVSSITASLRFDGALNVDLTEFQTNLVPYPRIHFPLATYAPVISAEKAYHEQLSVAEITNACFEPA
P0DPH8  ERPTYTNLNRLIGQIVSSITASLRFDGALNVDLTEFQTNLVPYPRIHFPLATYAPVISAEKAYHEQLSVAEITNACFEPA
P68366  ERPTYTNLNRLISQIVSSITASLRFDGALNVDLTEFQTNLVPYPRIHFPLATYAPVISAEKAYHEQLSVAEITNACFEPA

        300       310       320       330       340       350       360       370
Q6ZLZ9  NMMAKCDPRHGKYMACCLMYRGDVVPKDVNAAVATIKTKRTIQFVDWCPTGFKCGINYQPPTVVPGGDLAKVMRAVCMIS
Q8IFP3  SMMAKCDPRHGKYMACCLMYRGDVVPKDVNAAVATIKTKRSIQFVDWCPTGFKCGINYQPPTVVPGGDLAKVMRAVCMIS
Q71U36  NQMVKCDPRHGKYMACCLLYRGDVVPKDVNAAIATIKTKRTIQFVDWCPTGFKVGINYQPPTVVPGGDLAKVQRAVCMLS
P68363  NQMVKCDPRHGKYMACCLLYRGDVVPKDVNAAIATIKTKRSIQFVDWCPTGFKVGINYQPPTVVPGGDLAKVQRAVCMLS
P0DPH7  NQMVKCDPRHGKYMACCMLYRGDVVPKDVNAAIATIKTKRTIQFVDWCPTGFKVGINYQPPTVVPGGDLAKVQRAVCMLS
P0DPH8  NQMVKCDPRHGKYMACCMLYRGDVVPKDVNAAIATIKTKRTIQFVDWCPTGFKVGINYQPPTVVPGGDLAKVQRAVCMLS
P68366  NQMVKCDPRHGKYMACCLLYRGDVVPKDVNAAIAAIKTKRSIQFVDWCPTGFKVGINYQPPTVVPGGDLAKVQRAVCMLS

        380       390       400       410       420       430       440
Q6ZLZ9  NSTAIAEVFSRMDQKFDLMYAKRAFVHWYVGEGMEEGEFSEAREDLAALEKDYEEVGIESNEAEGEDEGYEADY
Q8IFP3  NSTAIAEVFSRMDQKFDLMYAKRAFVHWYVGEGMEEGEFSEAREDLAALEKDYEEVGIESNDGEGEDEGYE---
Q71U36  NTTAIAEAWARLDHKFDLMYAKRAFVHWYVGEGMEEGEFSEAREDMAALEKDYEEVGVDSVEGEGEEEGEE--Y
P68363  NTTAIAEAWARLDHKFDLMYAKRAFVHWYVGEGMEEGEFSEAREDMAALEKDYEEVGVDSVEGEGEEEGEE--Y
P0DPH7  NTTAIAEAWARLDHKFDLMYAKRAFVHWYVGEGMEEGEFSEAREDLAALEKDYEEVGVDSVEAEAE-EGEE--Y
P0DPH8  NTTAIAEAWARLDHKFDLMYAKRAFVHWYVGEGMEEGEFSEAREDLAALEKDYEEVGVDSVEAEAE-EGEE--Y
P68366  NTTAIAEAWARLDHKFDLMYAKRAFVHWYVGEGMEEGEFSEAREDMAALEKDYEEVGIDSYEDE-D-EGEE---
```

Binding Sites
- Pironetin
- Colchicine
- Vinblastine

**Supplementary Figure 5**: Multiple Sequence Alignment of α-tubulins including known drug binding sites. Multiple sequence alignment (MSA) of *P. falciparum* and human α-tubulin isoforms. Alignment was calculated with T-Coffee. Selected drug binding sites are highlighted: Pironetin (brown), Colchicine (green), Vinblastine (purple).
